# Mechanical unloading and exercise volume differentially modulate the gut microbiome and intestinal gene expression in mice

**DOI:** 10.64898/2026.11.11.765836

**Authors:** Seif Hundam, Mohammad Borhan Al-Zghoul, Shahed Ogiliat, Duha Makableh, Laith A. Abu-Haniyeh, Shadi Shahatit

**Affiliations:** Department of Basic Medical Veterinary Sciences, Faculty of Veterinary Medicine, Jordan University of Science and Technology, P.O. Box 3030, Irbid 22110, Jordan; Department of Biotechnology and Genetic Engineering, Faculty of Science and Art, Jordan University of Science and Technology, P.O. Box 3030, Irbid 22110, Jordan

**Keywords:** Hindlimb suspension, Exercise volume, Gut microbiome, Gut–muscle axis, Intestinal gene expression, Differential abundance, Oxidative stress

## Abstract

Mechanical unloading and physical activity can influence the gut microbiome and intestinal physiology, but the effects of exercise volume and prior hindlimb suspension remain incompletely understood. This study investigated how hindlimb suspension, different exercise volumes, and exercise following hindlimb suspension affect gut microbial composition and intestinal gene expression in mice. Mice were assigned to five groups: control (C), hindlimb suspension (HL), high-volume exercise (HE), low-volume exercise (LE), and hindlimb suspension followed by exercise (HLE). Cecal microbiota composition was characterized using 16S rRNA gene sequencing, followed by alpha- and beta-diversity analyses and LinDA differential-abundance analysis. Jejunal gene expression was assessed by quantitative PCR for markers related to oxidative stress, inflammation, metabolic signaling, and epithelial function. Hindlimb suspension reduced microbial richness, as indicated by the Observed, Chao1, and ACE indices, and was associated with a significant difference in overall microbial community composition compared with controls. It was also associated with reductions in Streptococcaceae/Streptococcus and Spirochaetota. Low-volume exercise produced the most extensive phylum-level differential-abundance findings, whereas high-volume exercise showed no significant phylum-level differences from controls. Both exercise groups (HE and LE) exhibited increased intestinal expression of GPX1, HMOX1, NFKB1, and SOD2. HE also increased IGF and MUC2 expression, whereas HLE reduced IGF expression compared with controls. Community composition did not differ significantly between HLE and HL, and LGR5 expression did not differ significantly among groups. In conclusion, hindlimb suspension was associated with reduced microbial richness and selective taxonomic changes, whereas low-volume exercise produced the most extensive phylum-level alterations in microbial composition. Exercise volume was also associated with distinct intestinal transcriptional responses, while exercise following mechanical unloading did not produce a microbial community composition significantly different from that observed following unloading alone. These findings highlight the context-dependent effects of mechanical unloading and exercise on gut microbial composition and intestinal gene expression.

## Introduction

“Muscle-gut axis” refers to the intricate communication network between skeletal muscle and the gut, and it has emerged as a critical determinant of systemic health. Skeletal muscle is not only a locomotor organ but also one of the body’s largest endocrine tissues, influencing distant organ systems through the release of myokines and metabolic signals (1). In parallel, the intestinal microbiome contributes to nutrient metabolism, immune homeostasis, and protection against intestinal pathogens (2,3).

Communication among these systems may involve microbial metabolites, host-derived signaling molecules, and immune-mediated pathways, underscoring the importance of understanding how changes in physical activity and musculoskeletal physiology affect the intestinal environment (4,5).

Physical exercise is an important modulator of the gut microbiome, although the magnitude and direction of microbial responses vary with exercise volume, intensity, duration, host characteristics, and experimental conditions (ref). Studies of athletes have reported differences in microbial diversity and functional potential compared with those of less active individuals, including changes in microbial pathways associated with carbohydrate metabolism and short-chain fatty acid production (6–8). Exercise-associated microbial changes have also been linked to shifts in the relative abundance of specific taxa, including Prevotella (9). However, the effects of exercise on microbial diversity and composition vary across studies, suggesting that exercise dose and physiological context are key determinants of the intestinal microbial response.

The effects on the intestine extend beyond microbial composition to include host transcriptional responses. Exercise alters metabolic demand and can influence oxidative stress-responsive and inflammatory pathways (10,11). These responses may reflect physiological adaptation, but their magnitude and biological consequences depend on the exercise protocol and the host’s condition (12). Antioxidant defense mechanisms, including superoxide dismutase and glutathione peroxidase, help regulate cellular redox status, while heme oxygenase-1 contributes to cellular stress responses (13). Nuclear factor kappa B-related signaling also helps regulate inflammatory and immune responses (14). Examining the expression of genes associated with these pathways can therefore provide insight into intestinal molecular responses to exercise.

The intestinal epithelium and its associated mucus layer provide complementary components of host–microbial interaction. Mucin 2, encoded by MUC2, is a major structural component of the intestinal mucus layer and contributes to the physical separation of luminal microorganisms from the epithelial surface (15). By contrast, LGR5 is commonly used as a marker of intestinal stem cells and is associated with epithelial renewal (16). Changes in the expression of these genes may provide information about selected aspects of the mucosal response to physiological interventions.

Hindlimb suspension is widely used to model the consequences of reduced weight-bearing activity and mechanical unloading in rodents. Previous studies have shown that simulated microgravity and hindlimb unloading can alter gut microbial composition and intestinal physiology (17–19). These findings suggest that unloading-associated changes may extend beyond skeletal muscle to involve the intestinal microbial ecosystem. However, the extent to which subsequent exercise modifies these responses remains incompletely understood.

Although exercise can influence the gut microbiome and intestinal molecular responses, greater exercise volume does not necessarily increase microbial diversity or yield a uniformly beneficial host response. Furthermore, the physiological effects of exercise performed after mechanical unloading may differ from those observed when exercise is performed without prior unloading. Therefore, further investigation is needed to determine whether low- and high-volume exercise elicit distinct microbial and intestinal transcriptional responses and whether exercise after hindlimb suspension reproduces the responses observed in non-suspended animals.

Accordingly, we investigated the effects of hindlimb suspension, low-volume exercise, high-volume exercise, and low-volume exercise following hindlimb suspension on the cecal microbiome and intestinal gene expression in mice. We characterized the microbial community composition using 16S rRNA gene sequencing and assessed genes associated with oxidative stress responses, inflammation, metabolic signaling, and intestinal epithelial biology by quantitative PCR, including Gpx1, Hmox1, Nfkb1, Sod2, Igf1, Muc2, and Lgr5. By integrating microbial community analyses with intestinal gene expression measurements, this study aimed to determine whether exercise volume and prior mechanical unloading were associated with distinct patterns of microbial and host responses. We hypothesized that these responses would vary by exercise volume and the physiological context in which exercise was performed.

## Materials and methods

All experimental procedures were conducted in accordance with institutional and national guidelines for the care and use of laboratory animals and were approved by the Institutional Animal Care and Use Committee of Jordan University of Science and Technology (JUST-ACUC; Approval number: 16/4/12/366).

### Study population and experimental design

Fifty 16-week-old male BALB/c mice were obtained from the animal house of Jordan University of Science and Technology. At the beginning of the experiment, the mice weighed approximately 25–30 g. Animals were randomly allocated to five experimental groups according to the experimental design: control (C; n = 20, high exercise (HE; n = 20), exercise following muscle atrophy (HLE; n = 10), high-level exercise group (HL; n = 10), and low exercise (LE; n = 20). The experimental groups and their respective interventions were maintained throughout the experimental period.

Mice were housed under controlled environmental conditions at approximately 25°C and 65% relative humidity, with a 16-h light/8-h dark cycle. Animals had ad libitum access to standard laboratory chow and tap water throughout the study. Mice were housed in cages at a density appropriate for the experimental housing conditions.

Exercise interventions were performed using an electric motorized rodent treadmill (PsymCon Model 35500, Lafayette Instrument, Lafayette, IN, USA). The exercise protocols were applied according to the predefined experimental design for each group.

### Induction of disuse muscle atrophy

Disuse muscle atrophy was induced using the established hindlimb-suspension model (20). After an initial period under normal weight-bearing conditions, mice assigned to the muscle-atrophy-related experimental groups underwent hindlimb suspension for two weeks.

For hindlimb suspension, mice were suspended by the tail using a specialized suspension apparatus such that the hindlimbs were prevented from bearing weight. At the same time, the animals retained unrestricted access to food and water and could move freely within the cage (21). Control animals remained under normal weight-bearing conditions throughout the corresponding experimental period.

### Exercise protocol

The exercise interventions were performed on a motorized treadmill. The low-exercise (LE) protocol consisted of four 10-min treadmill sessions per day at 15 m/min, with a 16-min rest interval between consecutive sessions, for a total of 28 exercise sessions per week. The high-exercise (HE) protocol consisted of the same treadmill sessions performed twice daily, with a 3-h interval between the two daily exercise periods, for a total of 56 exercise sessions per week.

For the HLE group, mice first underwent hindlimb suspension to induce disuse muscle atrophy and then completed the LE exercise protocol. The LE protocol was selected after hindlimb suspension because the higher exercise volume in the HE protocol was considered potentially stressful for mice recovering from disuse-induced muscle atrophy. Thus, the experimental design allowed evaluation of the response to low-volume exercise after muscle atrophy while avoiding the additional physiological stress of the high-volume exercise protocol.

### Sample collection

At the end of the experimental period, mice were euthanized under ketamine/xylazine anesthesia, and tissue samples were collected for histological, RT-qPCR, and microbiome analyses. For histological evaluation and RT-qPCR analysis, we obtained samples from six mice per group (n = 6/group) in the C, HE, HL, and LE groups, and from five mice (n = 5) in the HLE group. We collected the gastrocnemius muscle for histological assessment and the mid-jejunal region of the intestine for intestinal histology and RT-qPCR gene expression analysis.

For microbiome analysis, cecal contents were collected from a total of 50 mice, comprising 12 samples from C, 12 from HE, 9 from HL, 5 from HLE, and 12 from LE. Cecal contents were collected immediately after euthanasia and transferred into sterile tubes. Samples designated for molecular analyses were immediately snap-frozen in liquid nitrogen to minimize nucleic acid degradation and subsequently stored at −80 °C until downstream analyses in a CryoCube Series freezer (Eppendorf, Germany). Histological tissue samples were fixed in 10% neutral buffered formalin before further processing.

### Relative-Quantitative Real-Time PCR (RT-qPCR)

#### RNA Isolation and cDNA Synthesis

Mid-jejunal intestinal tissues were carefully dissected, immediately snap-frozen in liquid nitrogen, and stored at −80°C until RNA extraction. Tissue samples were homogenized in TRI Reagent® with a Bead Ruptor Elite-Bead Mill Homogenizer (OMNI International, Kennesaw, GA, USA).

Total RNA was extracted using the Direct-zol™ RNA MiniPrep kit (Zymo Research, CA, USA) according to the manufacturer’s instructions. RNA concentration and quality were assessed using a BioTek PowerWave XS2 spectrophotometer (BioTek Instruments, Inc., Winooski, VT, USA) and a Qubit 4 Fluorometer (Thermo Fisher Scientific, MA, USA). We also evaluated RNA integrity by agarose gel electrophoresis. For each sample, 500 ng of total RNA was reverse transcribed using PrimeScript™ RT Master Mix (Takara Bio Inc., Kusatsu, Japan) to generate complementary DNA (cDNA).

#### RT-qPCR reaction preparation and analysis

Quantitative real-time PCR was performed using a Rotor-Gene Q MDx 5-plex instrument (QIAGEN, Hilden, Germany) and Blastaq™ Green qPCR Master Mix (Applied Biological Materials Inc., Richmond, Canada).

Each 20-µL reaction contained 10 µL qPCR master mix, 2 µL forward primer (2 pmol), 2 µL reverse primer (2 pmol), 2 µL cDNA template, and 4 µL nuclease-free water. The amplification protocol consisted of an initial step at 50°C for 2 min, followed by denaturation at 95°C for 15 min and 40 amplification cycles of 95°C for 10 s, 57°C for 30 s, and 72°C for 10 s. A final melting-curve analysis was performed at 95°C for 20 s to confirm amplification specificity. Each cDNA sample was analyzed in technical duplicate. Relative gene expression was calculated using the comparative Ct method. Glyceraldehyde-3-phosphate dehydrogenase (GAPDH) was used as a reference gene for normalization. Primer specificity was confirmed by melting-curve analysis.

The nucleotide sequences of target genes were obtained from the National Center for Biotechnology Information (NCBI) Nucleotide database. Primers were designed using the PrimerQuest tool (Integrated DNA Technologies, IDT). Table 1 presents the complete primer sequences.

**Table 1:**
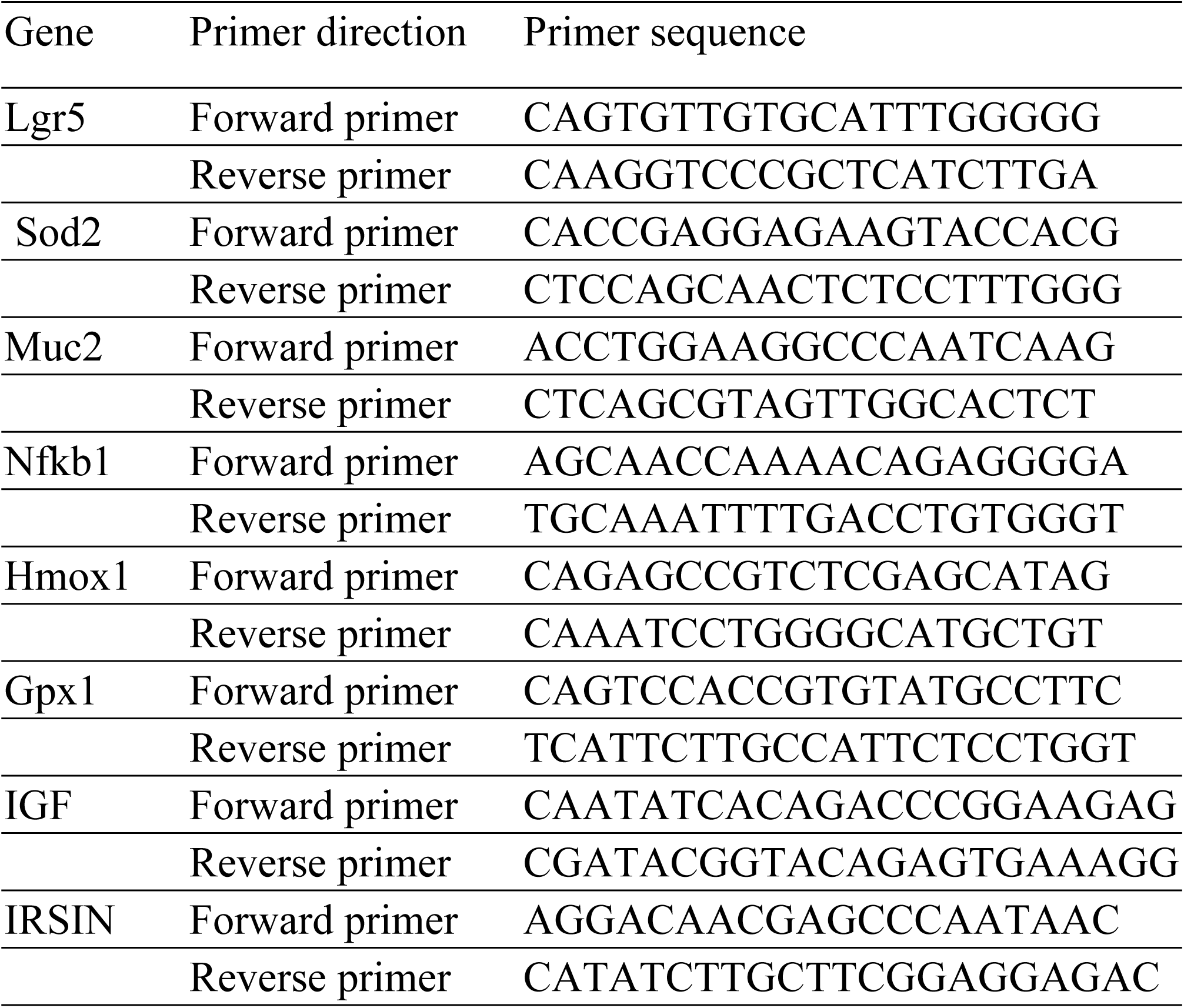
Primer sequences for RT-qPCR analysis.

### Microbiome Analysis

#### DNA Isolation and Sequencing

Cecal contents were collected aseptically from the experimental animals designated for microbiome analysis. We used approximately 150 mg of cecal content from each sample for microbial DNA extraction. Total microbial DNA was extracted using the DNeasy PowerSoil Pro Kit (QIAGEN, USA) according to the manufacturer’s instructions (QIAGEN, .June 2023.).

The bacterial community was characterized by sequencing the V3–V4 hypervariable regions of the 16S rRNA gene. We performed PCR amplification using the universal bacterial primers Bakt_341F (CCTACGGGNGGCWGCAG) and Bakt_805R (GACTACHVGGGTATCTAATCC). GenoDiscovery W.L.L. (Amman, Jordan) performed the sequencing commercially using a G99 platform (CA, USA), following the 2 × 300 bp paired-end sequencing protocol.

#### Data Analysis by Bioinformatics Tools

Raw 16S rRNA gene sequencing data were processed using QIIME 2 (version 2025.7) (22). We performed sequence quality control, denoising, and amplicon sequence variant (ASV) generation using DADA2. This workflow included quality filtering, removal of sequencing artifacts, and ASV inference. Taxonomic classification of ASVs was performed against the SILVA 138 reference database. The resulting feature table, taxonomy table, and sample metadata were subsequently imported into R for downstream statistical analysis.

For diversity analyses, the ASV feature table was rarefied to a common sequencing depth to minimize the influence of unequal sequencing depth among samples. The rarefied ASV table was used to calculate alpha- and beta-diversity metrics. Alpha diversity was assessed using Observed ASVs, Chao1, ACE, Shannon, and Simpson indices, and differences among the five experimental groups (C, HE, HL, HLE, and LE) were evaluated using the Kruskal–Wallis test followed by Dunn’s post-hoc test, with multiple comparisons adjusted using the Benjamini–Hochberg method. Beta diversity was evaluated using Bray–Curtis dissimilarity calculated from the rarefied community abundance table. We visualized community-level differences using principal coordinates analysis (PCoA), and assessed differences in microbial community composition among groups using PERMANOVA based on Bray–Curtis distances. Because within-group dispersion can influence PERMANOVA, we also performed PERMDISP to evaluate homogeneity of multivariate dispersion among experimental groups.

We performed differential-abundance analysis using LinDA (Linear models for Differential Abundance analysis), implemented in the MicrobiomeStat R package. LinDA was performed using the original non-rarefied ASV count table, independently at the Phylum, Family, and Genus levels. Six predefined biological contrasts were evaluated: HE vs. C, HE vs. LE, HLE vs. C, HLE vs. HL, HL vs. C, and LE vs. C. LinDA estimated log2 fold changes while accounting for the compositional nature of microbiome data. Positive log2 fold-change values indicated higher abundance in the second group of each contrast, whereas negative values indicated lower abundance. Multiple testing was controlled using the Benjamini–Hochberg procedure. Taxa with FDR < 0.05 were considered significantly differentially abundant. Volcano plots were generated separately for the Phylum, Family, and Genus levels across all six predefined contrasts.

## Results

### Alpha diversity across experimental groups

Alpha diversity analysis revealed differences in richness-related indices among the experimental groups, while diversity indices remained largely unchanged (Fig 1). For richness measures, including Observed, Chao1, and ACE, the HL group showed significantly lower richness than the control (p < 0.05), while the HE, HLE, and LE groups showed intermediate values that were not consistently different from either C or HL (p> 0.05). In contrast, diversity indices that account for both richness and evenness, including Shannon and Simpson, did not differ significantly among groups (p > 0.05).

**Fig 1.**
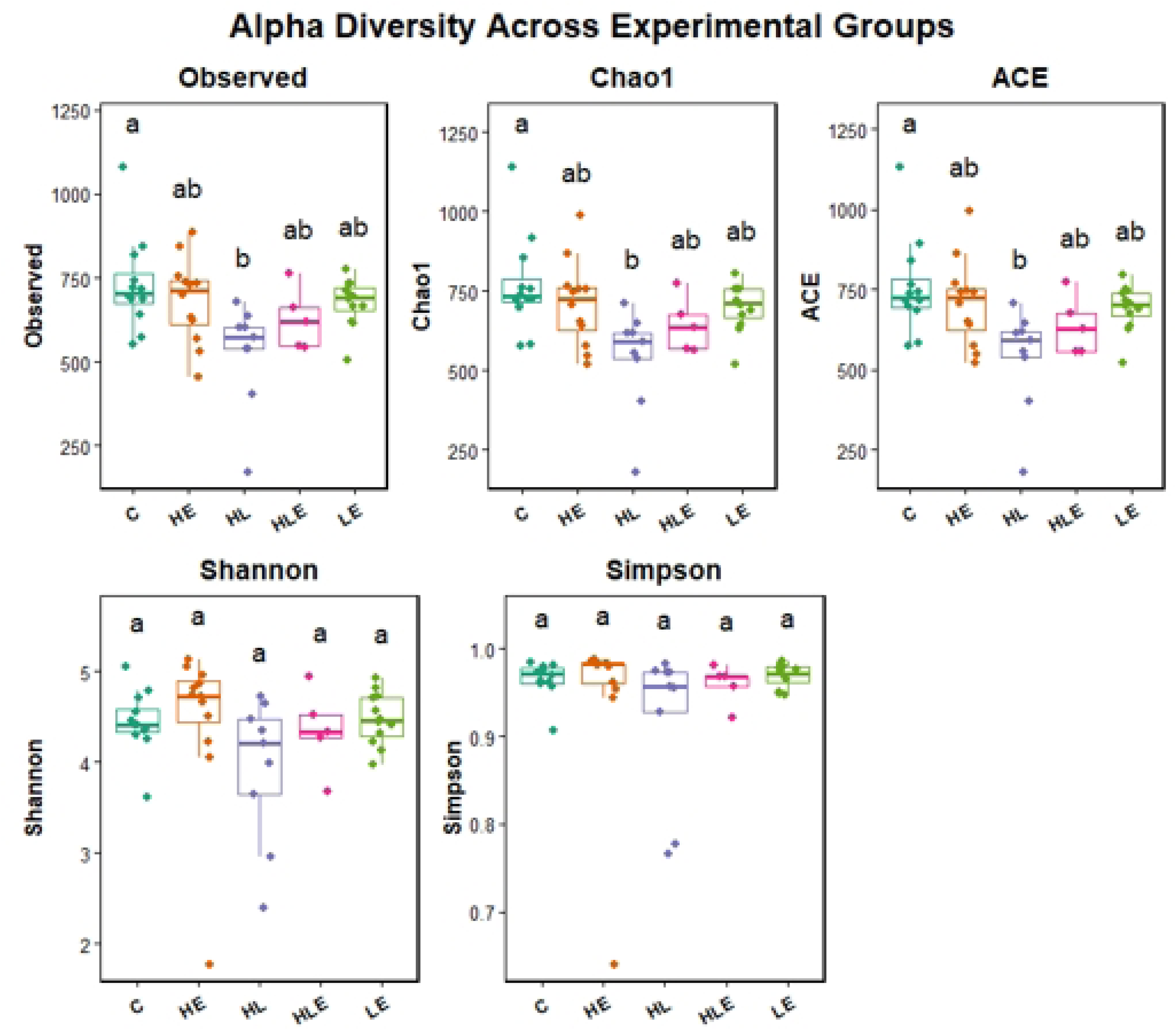
Boxplots showing alpha diversity indices, including Observed richness, Chao1, ACE, Shannon, and Simpson, across experimental groups: control (C), hindlimb suspension (HL), high-volume exercise (HE), low-volume exercise (LE), and hindlimb suspension followed by exercise (HLE). Richness indices (Observed, Chao1, and ACE) differed significantly among groups, while diversity indices (Shannon and Simpson) showed no significant variation. Different lowercase letters above the boxplots indicate statistically significant differences between groups (p < 0.05).

### Beta diversity across experimental groups

Bray–Curtis PCoA revealed a significant overall difference in microbial community composition among groups (PERMANOVA, R² = 0.119, p = 0.004; Fig 2). Pairwise analysis showed that the HL group significantly altered the microbiota compared to the control (p = 0.033). No significant differences were observed for the exercise rescue (HL vs. HLE; p = 0.16), exercise groups overall (p = 0.058), exercise intensity (HE vs. LE; p = 0.62), or rescue comparison (HL vs. HLE; p = 0.81).

**Fig 2.**
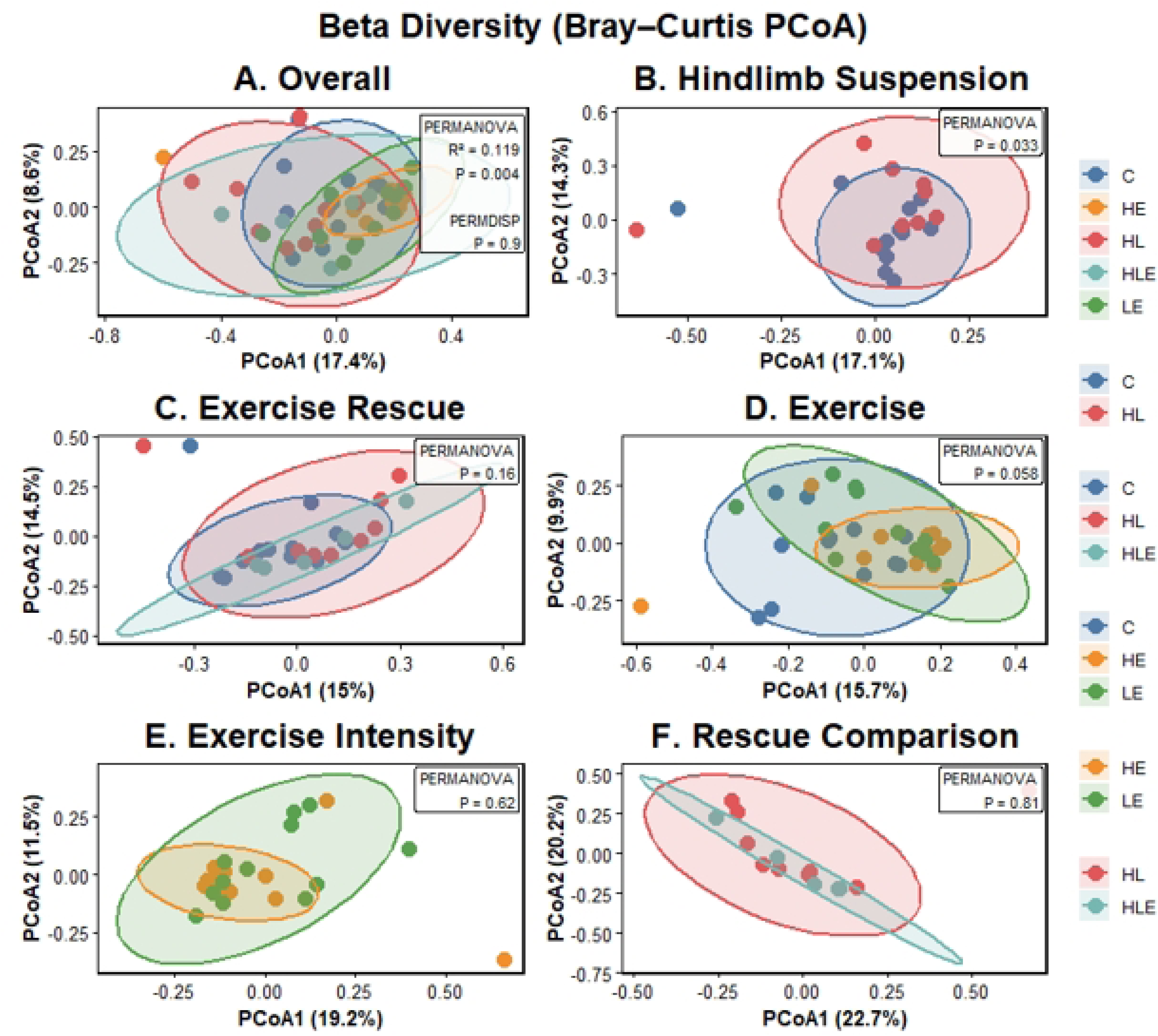
Beta diversity (Bray–Curtis PCoA). PCoA showing significant overall differences in microbial composition among groups (PERMANOVA, p = 0.004). HL differed significantly from C, while other comparisons were not significant. Ellipses represent 95% confidence intervals.

### Differential abundance analysis

At the phylum level (Fig 3), microbial communities were dominated by Firmicutes and Bacteroidota, with smaller proportions of other phyla across all groups. LinDA analysis at the phylum level revealed relatively few significant taxonomic changes among groups (Fig 4). The most pronounced differences were between the LE and control groups, with several phyla showing significant changes. Specifically, LE was associated with increased relative abundances of Bacteroidota (28.3% vs. 21.8%), Desulfobacterota (2.25% vs. 1.47%), and Verrucomicrobiota, as well as low-abundance taxa including Aenigmarchaeota. In comparison, Firmicutes showed a modest reduction (65.7% vs. 69.6%) compared with the control group. We also detected changes in rare taxa such as Actinobacteriota, Retaria, and Parabasalia.

**Fig 3.**
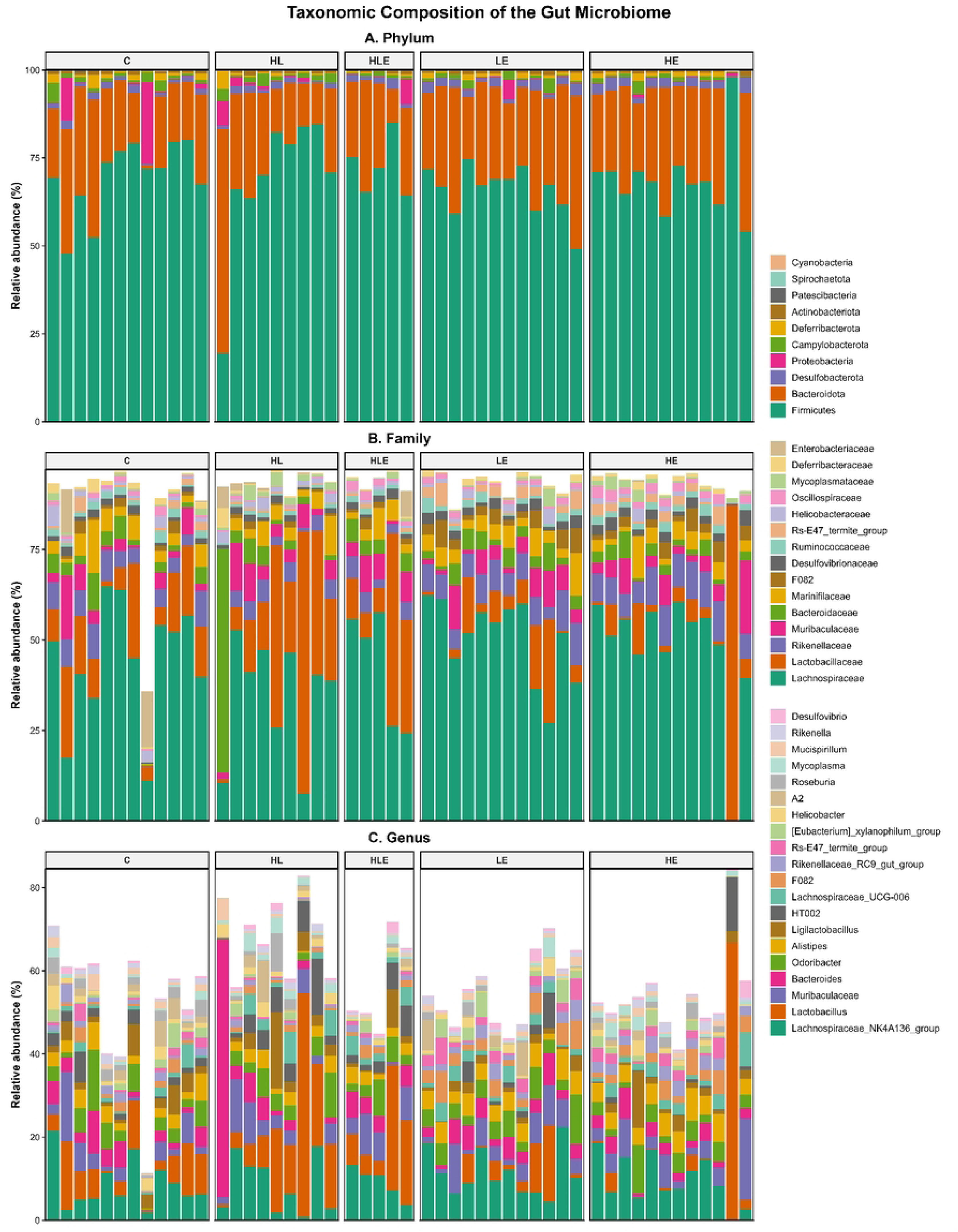
Taxonomic composition of the cecal microbiome. Relative abundance of bacterial taxa across control (C), hindlimb suspension (HL), hindlimb suspension followed by exercise (HLE), low-volume exercise (LE), and high-volume exercise (HE) groups at the (A) phylum, (B) family, and (C) genus levels. Each bar represents an individual sample, and taxa are shown according to their relative abundance within each sample. Only taxa represented in the respective panels are displayed.

**Fig 4.**
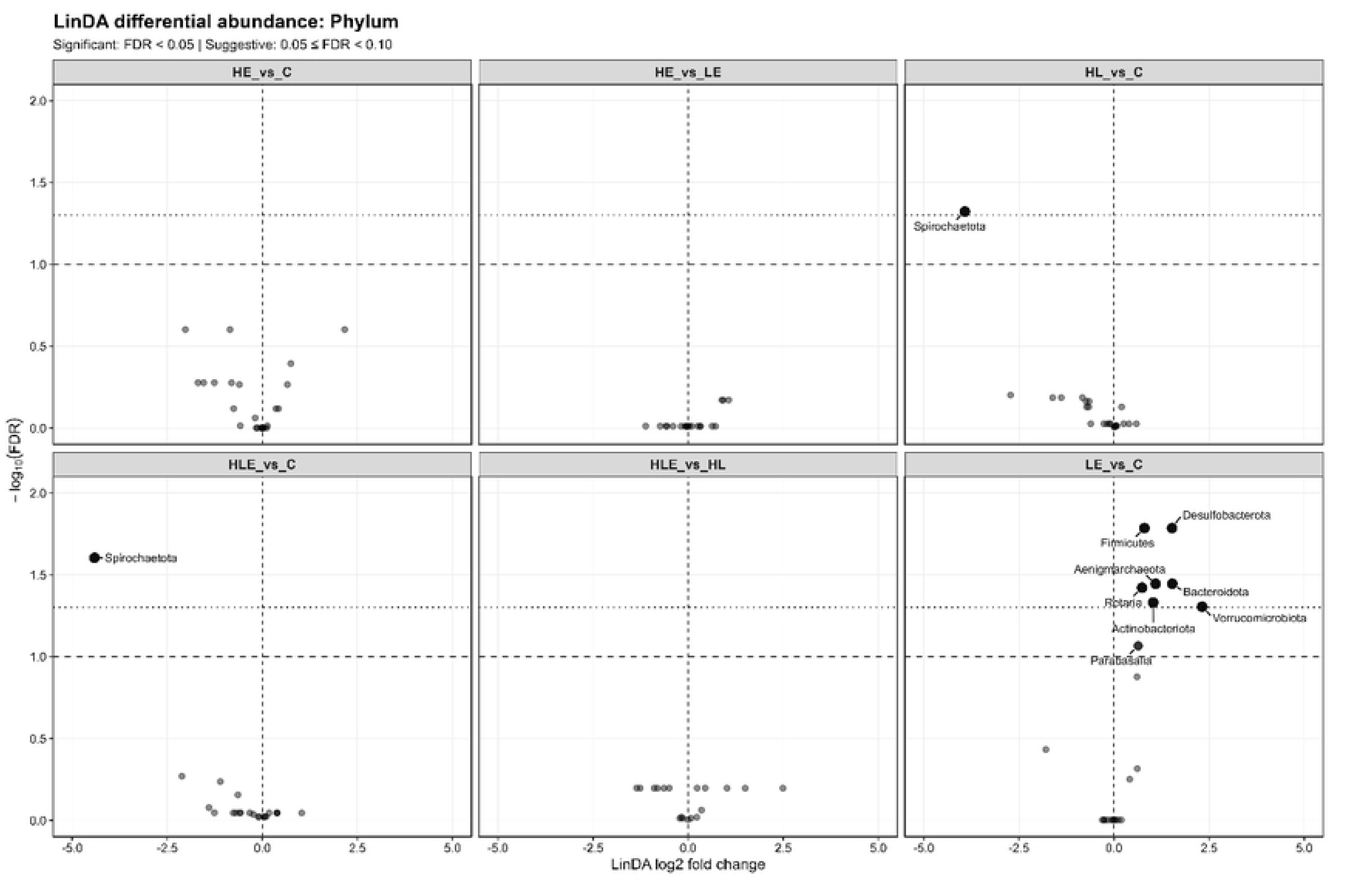
Differential abundance analysis at the phylum level using LinDA. Volcano plots showing differential abundance across pairwise comparisons: HE vs C, HE vs LE, HL vs C, HLE vs C, HLE vs HL, and LE vs C. The x-axis represents the LinDA log2 fold change, and the y-axis represents −log10(FDR). Taxa meeting the significance threshold (FDR < 0.05) are highlighted and labeled. Abbreviations: C, control; HL, hindlimb suspension; HLE, hindlimb suspension followed by exercise; LE, low-volume exercise; HE, high-volume exercise.

Comparisons involving hindlimb suspension showed fewer alterations. Spirochaetota was significantly reduced in both the HL and HLE groups relative to controls, with relative abundances of C: [0.058%], HL: [0.035%], HLE: [0.008%]. No significant phylum-level differences were detected between HE and C, HE and LE, or HLE and HL. Overall, these findings indicate that low-volume exercise induced the most pronounced phylum-level shifts in gut microbial composition, while hindlimb suspension exerted more limited effects.

At the family level, LinDA identified a significant reduction in Streptococcaceae in the HL group compared with controls (log2 fold change = −3.86, FDR = 0.002) (Fig 5). Consistent with this finding, the genus Streptococcus was also significantly reduced in HL compared with C (log2 fold change = −3.83, FDR = 0.004) (Fig 6). No other family- or genus-level taxa reached statistical significance in the examined pairwise comparisons.

**Fig 5.**
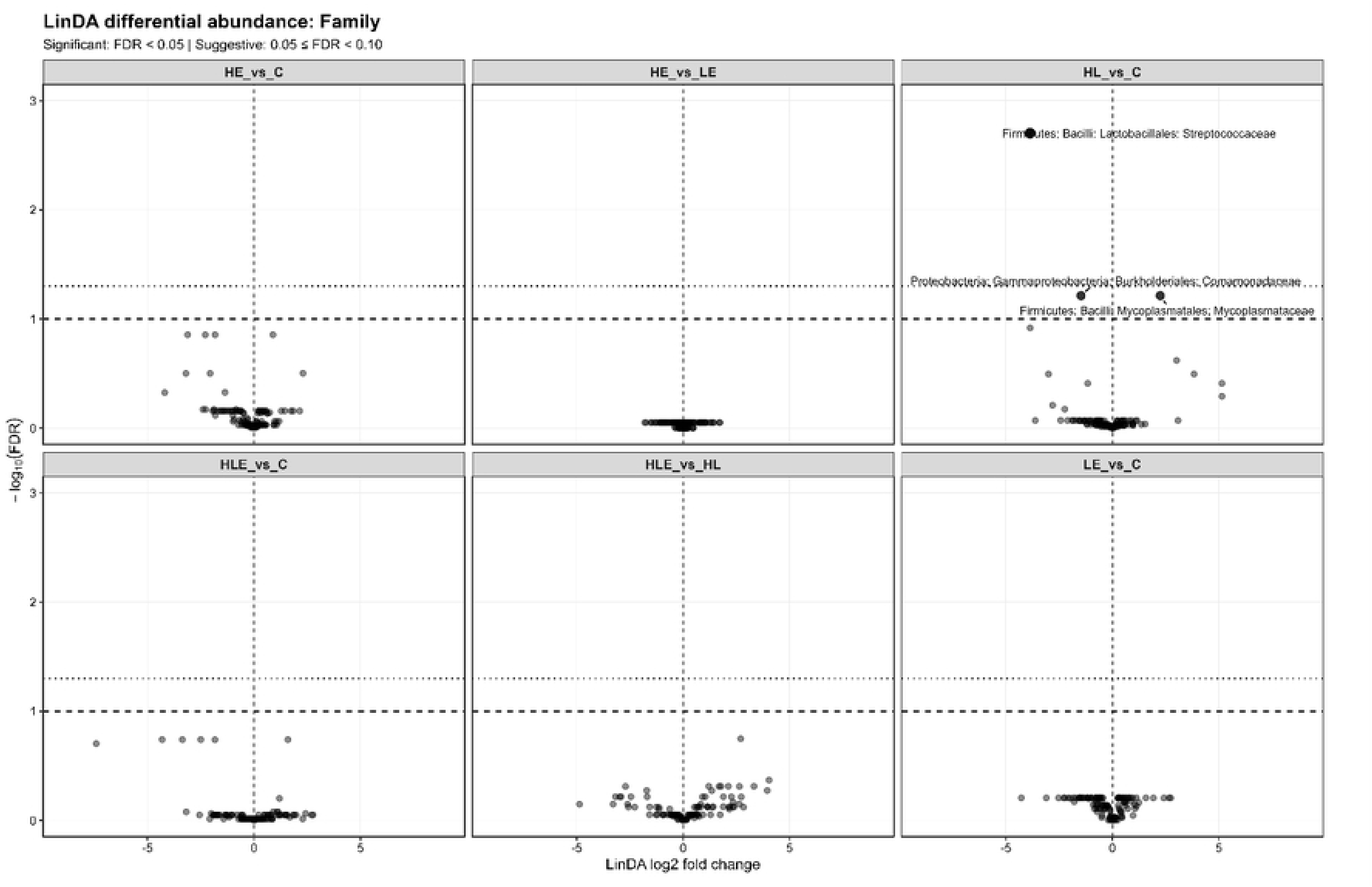
LinDA differential abundance analysis at the family level. Volcano plots showing differential abundance across pairwise comparisons: HE vs C, HE vs LE, HL vs C, HLE vs C, HLE vs HL, and LE vs C. The x-axis represents the LinDA log2 fold change, and the y-axis represents −log10(FDR). The dashed horizontal lines indicate the significance threshold (FDR < 0.05). Significant taxa are labeled.

**Fig 6.**
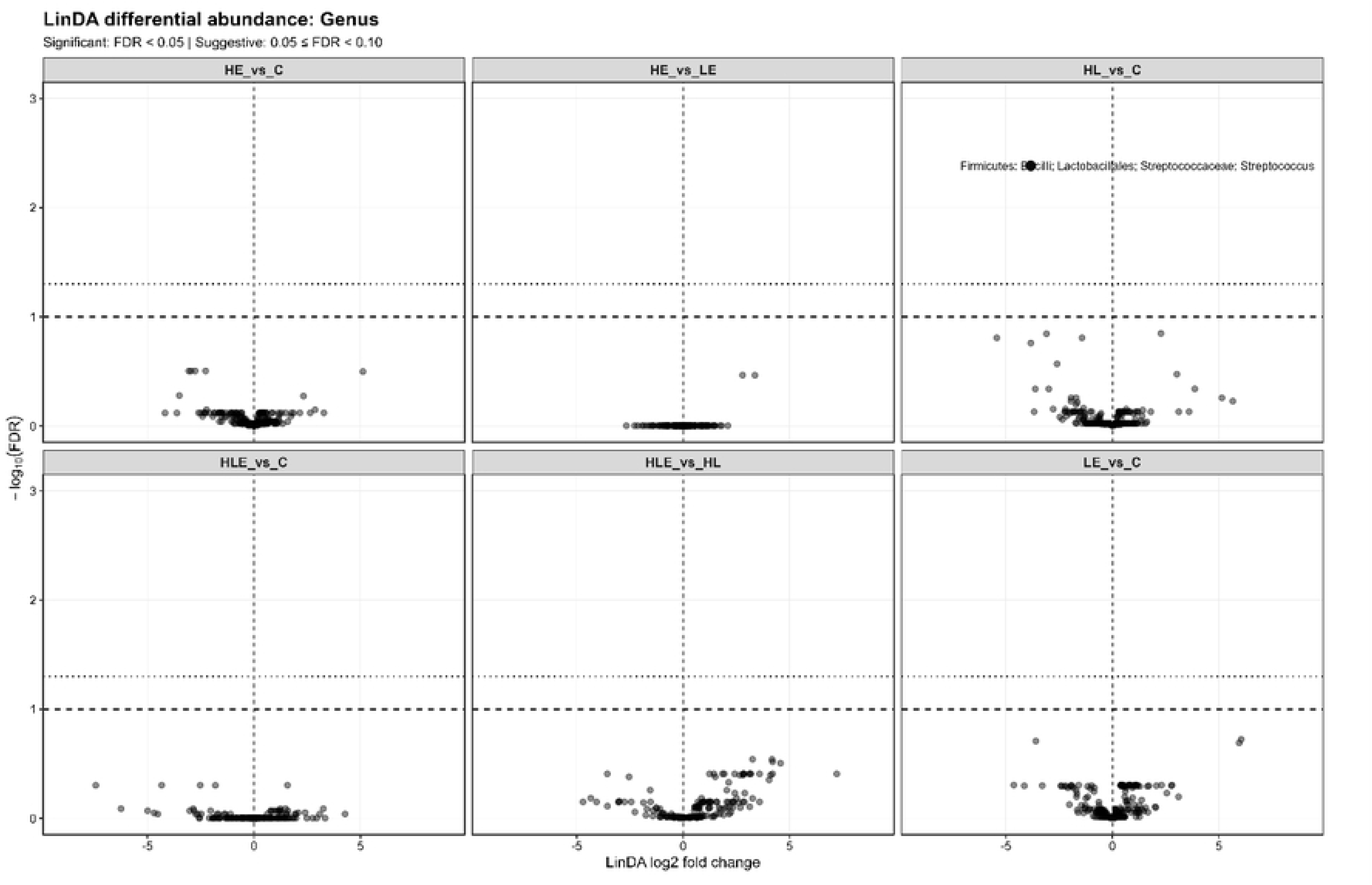
LinDA differential abundance analysis at the genus level. Volcano plots showing differential abundance across pairwise comparisons: HE vs C, HE vs LE, HL vs C, HLE vs C, HLE vs HL, and LE vs C. The x-axis represents the LinDA log2 fold change, and the y-axis represents −log10(FDR). The dashed horizontal lines indicate the significance threshold (FDR < 0.05). Significant taxa are labeled. Abbreviations: C, control; HL, hindlimb suspension; HLE, hindlimb suspension followed by exercise; LE, low-volume exercise; HE, high-volume exercise.

### Exercise modulates the expression of oxidative stress, inflammatory, and gut–muscle axis-related genes

qPCR analysis demonstrated treatment-specific changes in the expression of genes related to oxidative stress, inflammation, and metabolic signaling (Fig 7). GPX1 expression was significantly increased in the HE and LE groups compared with the control group, reaching approximately twofold higher expression, whereas the HL and HLE groups showed no significant differences. Similarly, HMOX1 expression was significantly elevated in both the HE and LE groups compared with the control.

**Fig 7.**
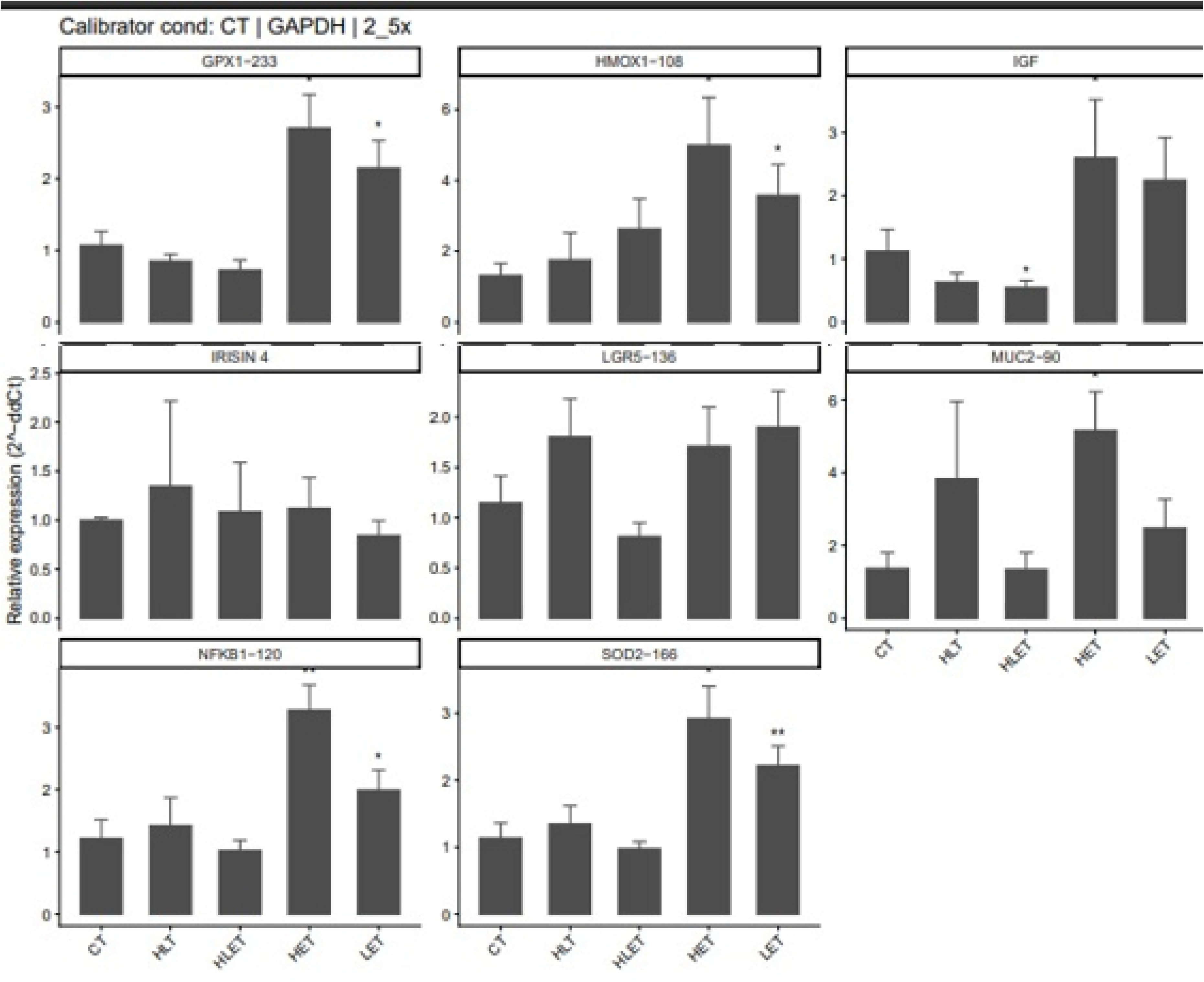
Relative expression of genes involved in oxidative stress, inflammation, metabolic signaling, and intestinal epithelial function. Relative mRNA expression of GPX1, HMOX1, IGF, IRISIN4, LGR5, MUC2, NFKB1, and SOD2 in control (C), hindlimb suspension (HL), hindlimb suspension followed by exercise (HLE), high-volume exercise (HE), and low-volume exercise (LE) groups. Expression was normalized to GAPDH and calculated relative to the control group using the 2^-ΔΔCt method. Bars represent mean ± SD. Statistical significance is indicated by *p < 0.05 and **p < 0.01.

IGF expression was significantly reduced in the HLE group compared with control, whereas expression was significantly increased in the HE group. The LE group showed a numerical increase in IGF expression compared with control, but this difference was not statistically significant. IRISIN4 expression did not differ significantly among groups.

For inflammatory and oxidative stress-related markers, NFKB1 and SOD2 expression were significantly increased in the HE and LE groups compared with control. Among intestinal epithelial markers, LGR5 expression did not differ significantly among groups, whereas MUC2 expression was significantly higher in the HE group compared with the other experimental groups.

Overall, the exercise interventions, particularly HE and LE, increased expression of oxidative stress- and inflammatory response-related genes, while HLE produced a distinct response characterized by reduced IGF expression.

## Disscussion

In the present study, hindlimb suspension clearly altered the gut microbial ecology, primarily by reducing microbial richness and shifting the overall community composition. The significant reductions in Observed, Chao1, and ACE richness in the HL group indicate fewer detected and estimated taxa following mechanical unloading, whereas the Shannon and Simpson indices remained unchanged. This pattern suggests that hindlimb suspension affected taxon richness more strongly than the overall distribution or evenness of the microbial community.

The significant PERMANOVA result further supports an unloading effect on community structure, as the HL microbiota differed significantly from that of control animals. These findings are consistent with previous studies showing that simulated microgravity and mechanical unloading can alter gut microbial composition and intestinal homeostasis (17,23–25). In particular, Qin et al (26) reported that mechanical unloading reduced ACE and Chao1 richness without significantly altering Shannon diversity, together with clear differences in β-diversity and shifts in bacterial taxa at multiple taxonomic levels.

Similarly, recent work using hindlimb unloading has suggested that recovery of microbial alterations may accompany improvements in intestinal barrier function and skeletal muscle outcomes, supporting a close relationship between unloading-induced physiological changes and the gut microbial ecosystem (27). Nevertheless, the relatively modest proportion of variation explained by treatment in the present study (R² = 0.119) indicates that hindlimb suspension contributed to the observed variation in microbial community composition rather than solely determining it. This finding suggests that factors beyond unloading may contribute to differences among animals.

The observed reduction in richness should also be interpreted in the context of the distinct physiological nature of hindlimb suspension. Previous systematic reviews report heterogeneous effects of exercise on gut microbial α-diversity, with some animal studies showing changes in richness or diversity, while others report minimal effects (28). Therefore, the reduced richness observed following hindlimb suspension should not be interpreted simply as a consequence of reduced physical activity. Unlike conventional sedentary conditions, hindlimb suspension produces pronounced mechanical unloading, reduced weight-bearing activity, and substantial alterations in musculoskeletal and metabolic physiology. These combined physiological changes may more strongly perturb the host–microbiome environment and may explain why the HL group showed a more pronounced richness phenotype than the exercise groups.

At the taxonomic level, LinDA analysis identified a pronounced reduction in Streptococcaceae and its genus Streptococcus in HL compared with controls, together with a significant reduction in the phylum Spirochaetota. The decrease in Streptococcaceae was particularly marked, with its relative abundance decreasing from approximately 1.60% in controls to 0.08% following hindlimb suspension. The concurrent reduction in Spirochaetota further suggests that mechanical unloading may alter specific microbial lineages that are sensitive to changes in the host physiological environment. However, the biological significance of these taxonomic shifts remains to be established. Because the present analysis used 16S rRNA gene sequencing, changes in relative abundance cannot directly establish alterations in microbial metabolic activity or function. Thus, the observed reductions in Streptococcaceae, Streptococcus, and Spirochaetota should be interpreted as compositional signatures of unloading rather than direct evidence of altered microbial function. Further metagenomic, metabolomic, or targeted functional analyses are needed to determine whether these taxonomic changes translate into meaningful alterations in microbial metabolic capacity.

Exercise produced a different pattern from hindlimb suspension. Interestingly, low-volume exercise produced the most significant phylum-level differences compared with control, including changes in Firmicutes, Bacteroidota, Desulfobacterota, Aenigmarchaeota, Retaria, Actinobacteriota, and Verrucomicrobiota. In contrast, HE versus C did not produce significant differential-abundance changes at the phylum level. This finding matters because it suggests the relationship between exercise volume and microbiome response was not simply linear. Therefore, the broader taxonomic response observed with LE in the present study may reflect a dose- and context-dependent microbial adaptation rather than a greater physiological stress response.

In the present study, Bacteroidota increased from 21.75% in controls to 28.34% in LE, while Desulfobacterota increased from 1.47% to 2.25%. Previous reviews have reported exercise-associated alterations in Bacteroidota/Bacteroidetes, although the direction and magnitude of this response vary substantially between studies (29–31). A recent systematic review and meta-analysis also concluded that exercise type and intensity can influence the balance between major bacterial phyla, highlighting the importance of exercise prescription when interpreting microbiome responses (30).

The finding that LE produced more significant microbial changes than HE deserves particular consideration. Previous literature does not support a simple model in which greater exercise intensity or volume invariably produces greater microbiome diversity or taxonomic change. Indeed, systematic reviews have reported highly variable microbiome responses across exercise intensities and intervention protocols (32,30). The present findings therefore suggest that the microbial response may follow a non-linear relationship with exercise dose. LE may have provided a sufficient physiological stimulus to alter the intestinal environment without producing the same host stress response associated with higher exercise volume. However, because we did not measure microbial metabolites, intestinal permeability, or functional microbial pathways, this interpretation remains speculative and should be tested in future studies.

The qPCR findings provide an important complementary perspective. Although HE produced relatively few significant taxonomic changes compared with control, it was associated with substantial changes in intestinal gene expression. Both HE and LE significantly increased GPX1 and HMOX1, and elevated NFKB1 and SOD2. This coordinated response suggests activation of antioxidant and stress-responsive pathways in response to exercise. Exercise increases metabolic demand and reactive oxygen species production, and transient increases in oxidative and inflammatory signaling can be part of the normal adaptive response rather than necessarily indicating pathological damage (10,11). The simultaneous elevation of SOD2 and GPX1 is consistent with activation of antioxidant defense mechanisms, while increased HMOX1 suggests activation of a cellular stress-response pathway (33–35). Thus, the transcriptional response observed here may represent an adaptive response to exercise-associated physiological stress.

The apparent dissociation between microbial and host transcriptional responses is also noteworthy. LE produced the most significant phylum-level microbial differences, whereas HE produced fewer significant taxonomic changes but still induced marked changes in GPX1, HMOX1, NFKB1, and SOD2. This suggests that broad microbial community changes and intestinal molecular responses are not necessarily associated. Previous systematic reviews similarly emphasize that exercise-associated microbiome responses are heterogeneous and that changes in microbial diversity do not consistently predict functional or metabolic outcomes (36,37). Consequently, these data support an association between exercise and both microbial and intestinal molecular responses but do not show that microbial changes directly caused the observed transcriptional alterations.

Intestinal IGF expression differed among the experimental groups, with a significant increase in the HE group and a significant decrease in the HLE group relative to the control group. In contrast, the LE group exhibited a numerical increase that did not reach statistical significance. These findings suggest that intestinal IGF transcription varies with exercise condition and the physiological context of exercise.

Dalise et al. (38) also examined the relationship between exercise dose and IGF-1 responses in rats, testing aerobic exercise and neuromuscular electrical stimulation. Their findings showed that exercise-associated IGF-1 responses differed by intervention and biological outcome measured. Hippocampal IGF-1 gene expression was associated with exercise performance in the aerobic exercise conditions, whereas circulating IGF-1 concentrations exhibited a different pattern across exercise doses.

In the present study, the significant increase in intestinal IGF expression in HE, together with the nonsignificant numerical increase in LE, is consistent with the possibility that exercise volume influences intestinal IGF transcription. However, the comparison with Dalise et al. should be interpreted cautiously because their study examined rats and focused on hippocampal gene expression and circulating biomarkers. In contrast, the present study assessed intestinal IGF mRNA in mice. Importantly, the present study assessed IGF mRNA expression and did not measure IGF protein abundance or downstream signaling activity. Therefore, the transcriptional changes cannot be directly interpreted as corresponding changes in IGF protein production or biological activity. Overall, these findings demonstrate that intestinal IGF expression varies across the experimental conditions, highlighting the importance of considering both exercise exposure and prior mechanical unloading when interpreting host gene-expression responses.

Intestinal epithelial markers responded differently to the experimental interventions. Muc2 expression was significantly higher in the HE group, whereas LGR5 expression did not differ significantly among the experimental groups. MUC2 encodes mucin 2, a major structural component of the intestinal mucus layer that contributes to the separation of luminal microorganisms from the epithelial surface (15). The increased MUC2 expression in the HE group may indicate a transcriptional response associated with mucus production under higher-volume exercise conditions. Exercise can influence intestinal physiology through changes in metabolic demand, splanchnic blood flow, and mucosal signaling; however, these effects depend on the exercise protocol and physiological context (39). In the present study, increased MUC2 expression was not accompanied by direct measurements of mucus thickness, mucin protein abundance, or intestinal permeability. Therefore, although the result is consistent with potential mucosal adaptation, it does not establish a thicker mucus layer or improved intestinal barrier function.

In contrast, LGR5 expression did not change significantly across experimental groups. LGR5 is commonly used as a marker of intestinal stem cells and is associated with intestinal epithelial maintenance and renewal (40). The lack of a significant change suggests that the interventions did not produce a detectable difference in Lgr5 transcription at the time of sampling. Nevertheless, this finding does not demonstrate that intestinal stem-cell activity or epithelial renewal was unchanged.

The contrasting responses of MUC2 and LGR5 are noteworthy. Whereas high-volume exercise increased expression of a gene involved in mucus production, it did not significantly change the stem-cell-associated marker. This pattern suggests that the mucosal response to exercise may be gene-specific. Moreover, the broader results show that HE was associated with increased MUC2 expression, along with changes in other intestinal genes, including GPX1, HMOX1, NFKB1, SOD2, and IGF. However, the concurrent changes do not establish a mechanistic relationship between these pathways or demonstrate that the observed microbial alterations caused the host transcriptional responses.

Overall, the mucosal gene-expression results identify increased MUC2 expression as a notable response to high-volume exercise, while LGR5 expression remained statistically unchanged. Further studies incorporating MUC2 protein measurements, mucus-layer assessment, epithelial histology, and intestinal permeability assays are needed to determine whether the transcriptional response corresponds to measurable changes in mucosal protection or barrier function.

Importantly, exercise following hindlimb suspension did not completely restore the gut microbiome toward the control phenotype. Spirochaetota remained significantly reduced in HLE compared with controls, and we detected no significant community-level difference between HL and HLE. This partial response aligns with the broader literature showing that exercise-induced microbiome changes depend heavily on the intervention protocol and physiological context (41,42). It also suggests that the biological consequences of mechanical unloading may persist even after exercise is reintroduced. The HLE-associated reduction in IGF expression further supports the idea that exercise after unloading represents a distinct physiological condition rather than simply a return to normal exercise physiology. Thus, these findings argue against interpreting HLE as a complete “rescue” intervention and instead support partial, context-dependent recovery.

An important strength of the present study is its integration of community-level microbiome analysis with taxon-specific differential abundance and intestinal gene-expression measurements. The alpha-diversity results indicate reduced richness after unloading, the beta-diversity analysis shows an overall treatment-associated shift in community structure, and LinDA identifies specific taxa that contribute to these differences. The qPCR results then demonstrate that exercise-associated changes at the host level can occur even when broad microbial community differences are limited. This multi-level response aligns with the literature emphasizing that exercise–microbiome interactions are complex and depend on exercise prescription, host condition, diet, experimental duration, and analytical methodology.

Several limitations should nevertheless be considered. First, 16S rRNA sequencing provides taxonomic rather than functional information; therefore, the observed changes cannot be directly linked to microbial metabolic activity or metabolite production. Second, the present study did not measure short-chain fatty acids, bile acids, intestinal permeability, tight-junction proteins, or circulating inflammatory mediators, limiting the ability to determine whether the microbial changes had functional consequences for intestinal physiology. Third, although the concurrent microbiome and qPCR findings support an association between exercise, microbial composition, and intestinal molecular responses, they do not establish directionality or causality. Future studies combining shotgun metagenomics or metatranscriptomics with metabolomics, intestinal barrier measurements, and muscle-specific molecular analyses would provide a stronger mechanistic assessment of the gut–muscle axis. Fecal microbiota transplantation or targeted microbial interventions would also be particularly valuable for determining whether exercise-associated microbial changes contribute directly to the recovery of muscle and intestinal function following unloading.

## Conclusion

In summary, these findings extend previous work by showing that the gut microbiome responds differently to mechanical unloading, exercise performed without prior unloading, and exercise introduced after unloading. Hindlimb suspension was associated with reduced microbial richness, altered community composition, and selective reductions in Streptococcaceae/Streptococcus and Spirochaetota. Low-volume exercise produced the broadest phylum-level taxonomic response, whereas both high- and low-volume exercise induced marked intestinal transcriptional responses in antioxidant and inflammatory pathways. Importantly, exercise after hindlimb suspension did not fully restore the microbial or molecular phenotype to control levels. These findings support a model in which exercise volume and prior mechanical unloading jointly determine the magnitude and direction of gut microbial and intestinal responses, providing further evidence for a context-dependent gut–muscle axis.

## Acknowledgments

The authors sincerely thank the Deanship of Research at Jordan University of Science and Technology for providing financial support for this research project. Grant numbers 526/2026 and 513/2026 were instrumental in facilitating this work. They also sincerely thank Eng. Amany Al-Rashadieh for invaluable insights and exceptional technical assistance.

## Data availability

All raw sequence data (FASTQ files) and QIIME 2 artifacts, including quality control files, feature tables, taxonomy tables, and rooted phylogenetic trees, associated with this study are publicly available. They can be accessed through the following link: https://github.com/saifhandam/Mice_Cecum.git (accessed on October 29, 2026). Additionally, the raw sequence data are available in the NCBI SRA database under accession PRJNA1552827.

## References

1. Saponaro F, Bertolini A, Baragatti R, Galfo L, Chiellini G, Saba A, et al. Myokines and Microbiota: New Perspectives in the Endocrine Muscle–Gut Axis. Nutrients. 2024;16(23):4032. doi:10.3390/nu16234032

2. Neish AS. Microbes in Gastrointestinal Health and Disease. Gastroenterology. 2009 Jan 1;136(1):65–80. doi:10.1053/j.gastro.2008.10.080

3. Kinross JM, Darzi AW, Nicholson JK. Gut microbiome-host interactions in health and disease. Genome Med. 2011 Mar 4;3(3):14. doi:10.1186/gm228

4. Rooks MG, Garrett WS. Gut microbiota, metabolites and host immunity. Nat Rev Immunol. 2016 Jun 1;16(6):341–52. doi:10.1038/nri.2016.42

5. Levy M, Blacher E, Elinav E. Microbiome, metabolites and host immunity. Host-Microbe Interact Bact. 2017 Feb 1;35:8–15. doi:10.1016/j.mib.2016.10.003

6. Clarke SF, Murphy EF, O’Sullivan O, Lucey AJ, Humphreys M, Hogan A, et al. Exercise and associated dietary extremes impact on gut microbial diversity. Gut. 2014 Dec 1;63(12):1913. doi:10.1136/gutjnl-2013-306541

7. Barton W, Penney NC, Cronin O, Garcia-Perez I, Molloy MG, Holmes E, et al. The microbiome of professional athletes differs from that of more sedentary subjects in composition and particularly at the functional metabolic level. Gut. 2018 Apr 1;67(4):625. doi:10.1136/gutjnl-2016-313627

8. Ticinesi A, Lauretani F, Tana C, Nouvenne A, Ridolo E, Meschi T. Exercise and immune system as modulators of intestinal microbiome: implications for the gut-muscle axis hypothesis. Exerc Immunol Rev. 2019 Feb 13;25.

9. Petersen LM, Bautista EJ, Nguyen H, Hanson BM, Chen L, Lek SH, et al. Community characteristics of the gut microbiomes of competitive cyclists. Microbiome. 2017 Aug 10;5(1):98. doi:10.1186/s40168-017-0320-4

10. Vargas-Mendoza N, Morales-González Á, Madrigal-Santillán EO, Madrigal-Bujaidar E, Álvarez-González I, García-Melo LF, et al. Antioxidant and adaptative response mediated by Nrf2 during physical exercise. Antioxidants. 2019;8(6):196.

11. He F, Li J, Liu Z, Chuang CC, Yang W, Zuo L. Redox mechanism of reactive oxygen species in exercise. Front Physiol. 2016;7:486.

12. Suzuki K, Tominaga T, Ruhee RT, Ma S. Characterization and Modulation of Systemic Inflammatory Response to Exhaustive Exercise in Relation to Oxidative Stress. Antioxidants. 2020;9(5):401. doi:10.3390/antiox9050401

13. He L, He T, Farrar S, Ji L, Liu T, Ma X. Antioxidants Maintain Cellular Redox Homeostasis by Elimination of Reactive Oxygen Species. Cell Physiol Biochem. 2017 Nov 17;44(2):532–53. doi:10.1159/000485089

14. Basak S, Hoffmann A. Crosstalk via the NF-κB signaling system. TNF Superfamily 2008. 2008 Jun 1;19(3):187–97. doi:10.1016/j.cytogfr.2008.04.005

15. Hansson GC. Mucins and the microbiome. Annu Rev Biochem. 2020;89(1):769–93.

16. van der Flier LG, Clevers H. Stem Cells, Self-Renewal, and Differentiation in the Intestinal Epithelium. Annual Review of Physiology. Annual Reviews; 2009. p. 241–60. 10.1146/annurev.physiol.010908.163145

17. Shi J, Wang Y, He J, Li P, Jin R, Wang K, et al. Intestinal microbiota contributes to colonic epithelial changes in simulated microgravity mouse model. FASEB J. 2017 Aug 1;31(8):3695–709. doi:10.1096/fj.201700034R

18. Jin M, Zhang H, Zhao K, Xu C, Shao D, Huang Q, et al. Responses of Intestinal Mucosal Barrier Functions of Rats to Simulated Weightlessness. Front Physiol [Internet]. 2018;Volume 9-2018. Available from: https://www.frontiersin.org/journals/physiology/articles/10.3389/fphys.2018.00729

19. Yuan L, Zhang R, Li X, Gao C, Hu X, Hussain S, et al. Long-term simulated microgravity alters gut microbiota and metabolome in mice. Front Microbiol [Internet]. 2023;Volume 14-2023. Available from: https://www.frontiersin.org/journals/microbiology/articles/10.3389/fmicb.2023.1100747

20. Marzuca-Nassr GN, Vitzel KF, Murata GM, Márquez JL, Curi R. Experimental model of hindlimb suspension-induced skeletal muscle atrophy in rodents. In: Pre-Clinical Models: Techniques and Protocols. Springer; 2018. p. 167–76.

21. Theilen NT, Jeremic N, Weber GJ, Tyagi SC. Exercise preconditioning diminishes skeletal muscle atrophy after hindlimb suspension in mice. J Appl Physiol. 2018;125(4):999–1010.

22. Bolyen E, Rideout JR, Dillon MR, Bokulich NA, Abnet CC, Al-Ghalith GA, et al. Reproducible, interactive, scalable and extensible microbiome data science using QIIME 2. Nat Biotechnol. 2019;37(8):852–7.

23. Siddiqui R, Qaisar R, Khan NA, Alharbi AM, Alfahemi H, Elmoselhi A. Effect of Microgravity on the Gut Microbiota Bacterial Composition in a Hindlimb Unloading Model. Life. 2022;12(11):1865. doi:10.3390/life12111865

24. Yuan L, Zhang R, Li X, Gao C, Hu X, Hussain S, et al. Long-term simulated microgravity alters gut microbiota and metabolome in mice. Front Microbiol [Internet]. 2023;Volume 14-2023. Available from: https://www.frontiersin.org/journals/microbiology/articles/10.3389/fmicb.2023.1100747

25. Tang H, Chai X, Qin X, Lian YE, Gao Y, Wang H, et al. The impact of mechanical unloading on the gut microbiota and the mitigating role of butyrate in bone loss. Int Immunopharmacol. 2025 Jun 26;159:114909. doi:10.1016/j.intimp.2025.114909

26. Qin X, Lian YE, Tang H, Chai X, Gao Y, Ma Y, et al. Lactobacillus rhamnosus GG mitigates bone loss induced by mechanical unloading via regulation of the gut-bone axis. Front Nutr [Internet]. 2026;Volume 12-2025. Available from: https://www.frontiersin.org/journals/nutrition/articles/10.3389/fnut.2025.1734220

27. Yu Y, Zheng Y, Zhang H, Fan X, Guo J, Sun L, et al. Abdominal LIPUS ameliorates simulated microgravity induced skeletal muscle atrophy via the gut-muscle axis. Npj Microgravity. 2025 Oct 27;11(1):73. doi:10.1038/s41526-025-00514-8

28. Cullen JMA, Shahzad S, Dhillon J. A systematic review on the effects of exercise on gut microbial diversity, taxonomic composition, and microbial metabolites: identifying research gaps and future directions. Front Physiol [Internet]. 2023;Volume 14-2023. Available from: https://www.frontiersin.org/journals/physiology/articles/10.3389/fphys.2023.1292673

29. Motiani KK, Collado MC, Eskelinen J joonas, Virtanen KA, Löyttyniemi E, Salminen S, et al. Exercise training modulates gut microbiota profile and improves endotoxemia. Med Sci Sports Exerc. 2019;52(1):94.

30. Min L, Ablitip A, Wang R, Luciana T, Wei M, Ma X. Effects of exercise on gut microbiota of adults: a systematic review and meta-analysis. Nutrients. 2024;16(7):1070.

31. Gao R, Wang J, Song J, Zhang W, Quan L. Acute high-intensity exercise alters gut microbiota composition and energy metabolism in different strains of mice. Front Microbiol. 2026;17:1790697.

32. Cullen JM, Shahzad S, Dhillon J. A systematic review on the effects of exercise on gut microbial diversity, taxonomic composition, and microbial metabolites: identifying research gaps and future directions. Front Physiol. 2023;14:1292673.

33. Li S, Yan T, Yang JQ, Oberley TD, Oberley LW. The role of cellular glutathione peroxidase redox regulation in the suppression of tumor cell growth by manganese superoxide dismutase. Cancer Res. 2000;60(14):3927–39.

34. Ekoue DN, He C, Diamond AM, Bonini MG. Manganese superoxide dismutase and glutathione peroxidase-1 contribute to the rise and fall of mitochondrial reactive oxygen species which drive oncogenesis. Biochim Biophys Acta BBA-Bioenerg. 2017;1858(8):628–32.

35. Ryter SW. Heme oxygenase-1: an anti-inflammatory effector in cardiovascular, lung, and related metabolic disorders. Antioxidants. 2022;11(3):555.

36. Munukka E, Ahtiainen JP, Puigbó P, Jalkanen S, Pahkala K, Keskitalo A, et al. Six-week endurance exercise alters gut metagenome that is not reflected in systemic metabolism in over-weight women. Front Microbiol. 2018;9:2323.

37. Cullen JM, Shahzad S, Dhillon J. A systematic review on the effects of exercise on gut microbial diversity, taxonomic composition, and microbial metabolites: identifying research gaps and future directions. Front Physiol. 2023;14:1292673.

38. Dalise S, Cavalli L, Ghuman H, Wahlberg B, Gerwig M, Chisari C, et al. Biological effects of dosing aerobic exercise and neuromuscular electrical stimulation in rats. Sci Rep. 2017;7(1):10830.

39. van Wijck K, Lenaerts K, Grootjans J, Wijnands KA, Poeze M, Van Loon LJ, et al. Physiology and pathophysiology of splanchnic hypoperfusion and intestinal injury during exercise: strategies for evaluation and prevention. Am J Physiol-Gastrointest Liver Physiol. 2012.

40. Barker N, Van Es JH, Kuipers J, Kujala P, Van Den Born M, Cozijnsen M, et al. Identification of stem cells in small intestine and colon by marker gene Lgr5. Nature. 2007;449(7165):1003–7.

41. Finderle J, Schleicher VS, Schleicher LMS, Krsek A, Braut T, Baticic L. Exercise-induced modulation of the gut microbiota: mechanisms, evidence, and implications for athlete health. Gastrointest Disord. 2025;8(1):1.

42. Reljic D, Hermann HJ, Dieterich W, Neurath MF, Zopf Y. Exercise improves gut microbial metabolites in an intensity-dependent manner: a pooled analysis of randomized controlled trials. Gut Microbes. 2025;17(1):2579354.

